# Thermal fluctuations assist mechanical signal propagation in coiled-coil proteins

**DOI:** 10.1101/2020.05.04.076919

**Authors:** Judit Clopés, Jaeoh Shin, Marcus Jahnel, Stephan W. Grill, Vasily Zaburdaev

## Abstract

Recently it has been shown that the long coiled-coil membrane tether protein Early Endosome Antigen 1 (EEA1) switches from a rigid to a flexible conformation upon binding of a signaling protein to its free end. This flexibility switch represents a novel motor-like activity, allowing EEA1 to generate a force that moves vesicles closer to the membrane they will fuse with. To elucidate how binding of a single signaling protein can globally change the stiffness of a 200 nm long chain, we propose a simplified description of the coiled-coil as a one-dimensional Frenkel-Kontorova chain. Using numerical simulations, we find that an initial perturbation of the chain can propagate along its whole length in the presence of thermal fluctuations, changing the configuration of the entire molecule and thereby affecting its stiffness. Our work sheds light onto intramolecular communication and force generation in long coiled-coil proteins.

---

Coiled-coils (CC) are structural protein motifs composed of two or more intertwined *α*-helices. They play essential roles in multiple cellular processes, including gene regulation, muscle contraction, and cell signaling [1]. Early Endosome Antigen 1 (EEA1) is an extended fibrous protein with a large dimeric CC domain that guides vesicles coated with the small GTPase Rab5 to early endosomes [2]. It has recently been shown that unbound EEA1 is rather stiff and becomes more flexible when active Rab5 selectively binds to its N-terminus [3]. The entropic force of the flexible chain brings the tethered vesicle closer to the endosome membrane to facilitate the downstream vesicle fusion. Next, GTP hydrolysis breaks the interaction between EEA1 and Rab5, suggesting reversibility of this process. Similar behavior has been observed for other long CC tethering proteins, such as the golgin family [4–7]. However, the underlying mechanism of the conformational switches is not fully understood. In particular, how does a local perturbation induced by a ligand binding at one end of the CC propagate through the whole structure of the protein to change its global flexibility?

Structurally, the center part of EEA1 folds as a long canonical homodimeric CC of approximately 200 nm (1275 amino acids) on almost its total length [8]. The sequence of *α*-helices forming ideal left-handed CCs shows a periodic structural motif of *hpphppp* known as heptad repeat, where *h* and *p* are the hydrophobic and polar residues, respectively [9]. In an aqueous environment, the CC fold is formed by the assembly of *α*-helices driven mainly by hydrophobic interactions established between the *h* residues of different strands. The regularity in the *α*-helix sequence is interrupted by short discontinuity regions distributed along the structure (see Fig. 1, (a)). While the nature of the fold in these regions is not known, the recent findings have related the flexibility change with the presence of such discontinuities; the mutant EEA1 that lacks these regions does not show the stiffness switch [3]. Therefore, we hypothesize that the CC regions could be involved in the energy transmission of the Rab5-EEA1 interaction through the whole structure whereas the discontinuities could be responsible for inducing the conformational change. To ensure energy transmission, the ideal CC regions in EEA1 (without discontinuities) must have a structure capable of propagating the perturbation with a sufficient signal amplitude to the discontinuity regions without destroying the dimeric CC fold.

**FIG. 1.**
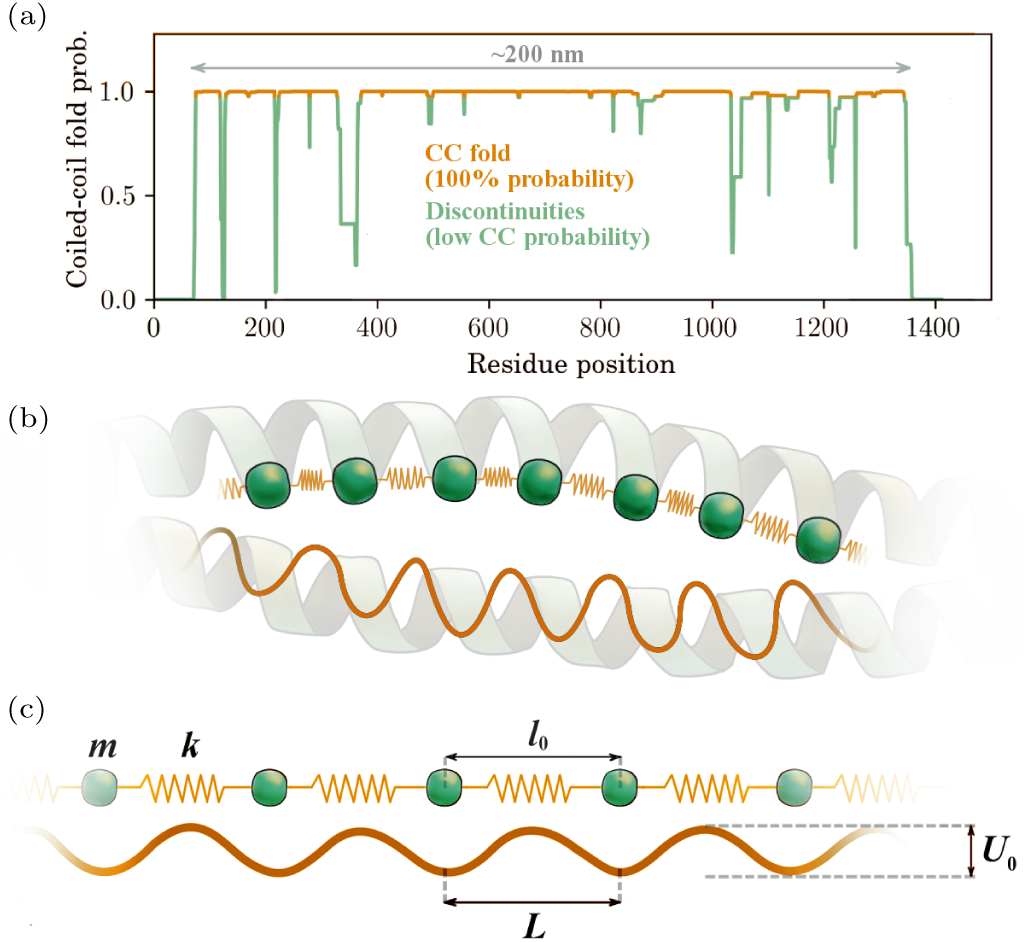
Modeling hydrophobic interactions between strands in a canonical coiled-coil structure (a) Coiled-coil fold prediction for EEA1 in humans. Prediction plot computed with PCOILS bioinformatics toolkit [11] using a window of 28 amino acids. (b) Sketch of the application of the Frenkel-Kontorova model to map the hydrophobic contacts stabilizing the two *α*-helical strands forming a homodimeric canonical coiled-coil protein. (c) Frenkel-Kontorova model represented by a chain of beads connected by harmonic springs in the periodic potential. *U*_0_ and *L* are the strength and the period of the hydrophobic interaction potential respectively, *l*_0_ is the half turn separating two hydrophobic residues and the equilibrium length of the harmonic spring connecting two beads with the spring constant *k*. Note that *l*_0_ coincides with *L* as both *α*-helices in EEA1 are identical in the amino acid sequence.

The main interest of this work lies in understanding the feasibility of this energy transmission mechanism from the physical point of view. For that, we applied an over-damped one-dimensional Frenkel-Kontorova (FK) model to a novel range of problems of signal transduction. We analyzed the dynamics of dislocations in FK chains for the length and time scales characteristic to the EEA1 protein, and we find that thermal fluctuations are necessary for the signal to propagate through the whole structure. Moreover, we provide the range of parameters when such signal propagation is feasible within the experimentally measured time. Finally, we briefly discuss how the small perturbation can trigger the change of the global properties of EEA1 in the framework of our model.

## a. The Frenkel-Kontorova model applied to coiled-coil structures

The hydrophobic interactions between strands in an ideal CC fold can be described in terms of a one-dimensional FK chain [10, 12] (see Fig. 1, (b) and (c)). The original FK model was introduced in the late 1930s to describe defect dynamics in solids. In the context of biological systems, it has been used to model phenomena such as friction [13, 14], protein folding [15], chromatin dynamics [16], and DNA breathing [17]. When applied to CCs, this model is coarse-grained to the hydrophobic residue level, where each heptad repeat is described by two beads connected by a spring. The periodic potential with amplitude *U*_0_ and period *L* characterizes the strength of the hydrophobic interaction between residues of opposite strands. Since the two *α*-helices forming the dimeric CC in EEA1 have an identical amino acid sequence and align in parallel [8], the effect of the hydrophobic interactions in the FK chain was chosen to be commensurate where the equilibrium length of the springs *l*_0_ coinsides with the potential period *l*_0_ = *L*. The elastic constant *k* of the spring connecting two neighboring beads is related to the contributions from both the rigidity at small stretches and the hydrogen bonds established between the 3rd and 4th nearest-neighbors in *α*-helices (half heptad repeat). We have modeled the canonical coiled-coil regions in EEA1 by means of an overdamped FK chain. We first show that thermal fluctuations are necessary to ensure the propagation of an initial perturbation on one extreme through the whole length of the chain. Then, we study the chain parameters which allow for the signal transmission and relate these results to the EEA1 protein.

The equation of motion for the *i* -th bead in an over-damped FK chain in terms of Langevin dynamics is

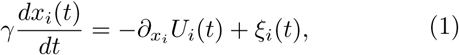

where *x*_*i*_(*t*) is the position of the bead, *γ* is the friction introduced by both the medium and the interaction between opposite strands, and *ξ*_*i*_(*t*) is a Gaussian white noise describing thermal fluctuations, satisfying ⟨*ξ*_*i*_(*t*)⟩ = 0 and ⟨*ξ*_*i*_(*t*)*ξ*_*j*_(*t*′)⟩ = 2*γk*_*B*_*Tδ*(*t* − *t*′)*δ*_*i,j*_. The potential energy *U*_*i*_(*t*) is given as

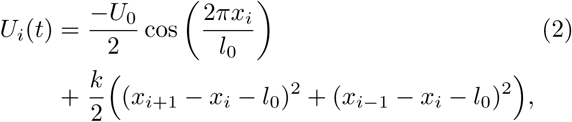

In the absence of thermal noise, the compression or expansion (kink and antikink) introduced between a pair of particles in an overdamped commensurate FK chain relax to a finite length characterized by 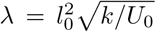 [12, 13]. This length λ, known as kink width, determines the region of the chain affected by a compression or an expansion of length *l*_0_.

We introduce the Rab5-EEA1 interaction as a register shift leading to a relative sliding of half a heptad repeat in one strand with respect to the opposite. This mechanism could introduce long distance communication along the CC fold at a low energetic cost, since it does not involve the exposure of hydrophobic residues to the aqueous environment. A similar sliding mechanism has been suggested to be responsible for the allosteric communication along the homodimeric CC forming the stalk of dynein [19, 20], and is also involved in signal transmission along transmembrane proteins [21]. In our model, the initial perturbation from Rab5 binding is introduced by generating a compression (kink) of length *l*_0_ between the leftmost pair of beads of the FK chain and keeping the position of the first bead fixed. We then analyzed the dynamics of the potential energy distribution per bead on the FK chain *U*_*i*_(*t*). The signal propagation analysis was performed by averaging single kink trajectories for kinks introduced in one of the extremes and in the center of the chain. The signals were post-processed using the Savitzky-Golay filter [18]. We then analyzed the propagation front *x*_fr_ (see Fig. 2 (c)), defined as the length at which the filtered signal amplitude reaches the energy offset, which is the mean energy of the beads introduced solely by temperature in a resting chain.

**FIG. 2.**
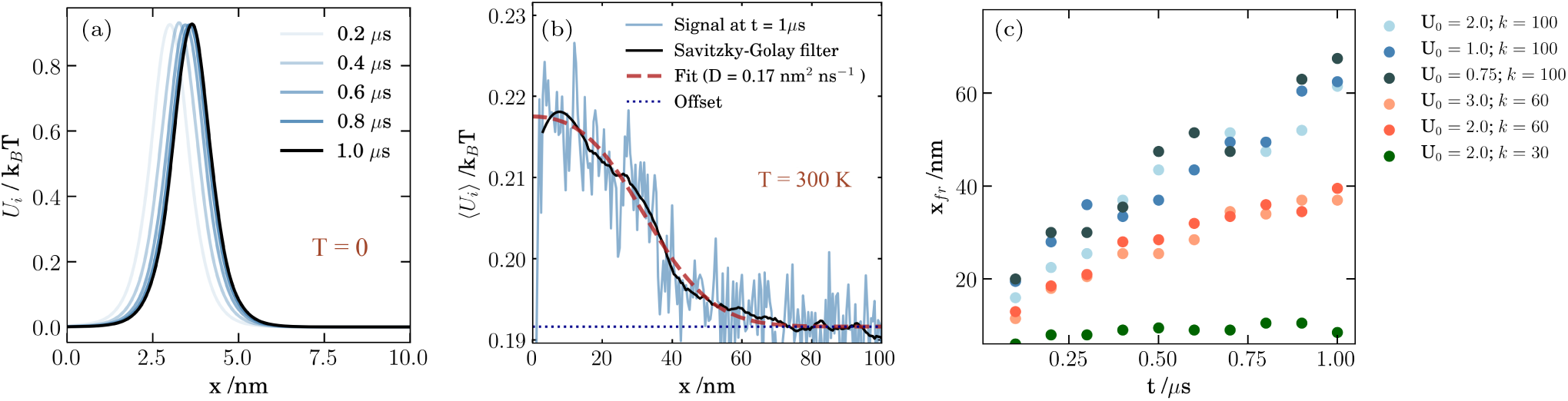
Signal propagation analysis. (a) Evolution of the potential energy distribution profile along a Frenkel-Kontorova chain at zero temperature with *U*_0_=1 *k*_*B*_ *T* and k=100 *k*_*B*_ *T* nm^−2^. The initial contraction relaxes and does not propagate. (b) Potential energy distribution profiles averaged over 5000 trajectories for identical chains as in (a), but now affected by thermal fluctuations. In this case, the compression is active and propagates diffusively. The averaged signal has been filtered by applying the Savitzky-Golay algorithm with a window size of 51 data points [18]. The fit curve (maroon dashed line) corresponds to the solution to the diffusion equation with a reflecting boundary at *x* = 0 with the diffusion constant *D* extracted from the fit. (c) Dynamics of the position of the potential energy front *x*_fr_ for different *U*_0_ and *k* pairs. The front position *x*_fr_ is the position where the filtered signal reaches the offset. The units of *U*_0_ and *k* are *k*_*B*_ *T* and *k*_*B*_ *T* nm^−2^ in all plots, respectively.

## b. Characterization of the signal transmission

The energy transport along the FK chain is a result of two processes: the relaxation of the contraction and a diffusive propagation of the kink due to thermal effects. In the absence of thermal fluctuations, or at low temperature, the contraction applied to the extreme relaxes to a width determined by λ, and does not propagate further (see Fig. 2, (a)). At intermediate temperatures (*U*_*i*_ ⪆ *k*_*B*_*T*), a single kink diffuses while maintaining a constant width λ. Higher temperatures (*U*_*i*_ < *k*_*B*_*T*) melt the coiled-coil structure by spontaneously generating kink-antikink pairs along the chain. The relaxation of the kink at zero temperature occurs much faster than the characteristic diffusion time. Therefore, both phenomena can be decoupled [22], and the diffusive regime is recovered at large time scales (see Fig. 2, (b)).

In the absence of thermal fluctuations, the chain remains on its new energy minimum configuration without any motion once the elastic energy accumulated by the register shift 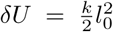 is released. Thermal fluctuations enable the chain monomers to cross the potential barrier yielding further propagation of the energy. The kink jumps are, therefore, more frequent as the potential depth *U*_0_ decreases, so the overall signal propagation is faster for decreasing *U*_0_. Furthermore, the propagation is enhanced in stiffer chains, i.e., by increasing the elastic constant *k*, since the energy introduced in the initial perturbation is greater (see Fig. 2 (c)). This phenomenon is reminiscent of the motion of a Brownian particle or a polymer in a tilted washboard potential [23]. The register shift in our model plays a similar role of tilting the potential.

The potential energy distribution at chain positions away from the boundary *x* > λ shows a diffusive evolution at large times (see Fig. 2 (b)). The fit (maroon dashed line) results from the solution of the diffusion equation with a reflecting boundary at *x* = 0 and with the initial condition as the Dirac delta function at *x*_pk_.

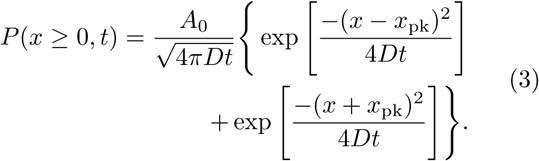

where the diffusion constant *D* is extracted from the fit and *A*_0_ is the normalization constant. In our situation, *x*_pk_ is expected to be very close to reflecting boundary, at the distance of the order of the width of the kink. For simplicity, we do not specify this value and just use it as an additional fit parameter.

## c. Relation to the case of the EEA1 coiled-coil protein

The parameter set used to reproduce the physiological conditions at which CC proteins are functional is listed in Table I. The equilibrium distance between two beads corresponds to the length of a turn in canonical CCs, *l*_0_, which has an approximate value of 0.5 nm [24, 25]. The range of the elastic constant *k* corresponds to the stretching of coiled-coil regions of myosin in the limit of small pulling forces, where the elasticity is entropic, until the first hydrogen bond in the *α*-helix (half heptad repeat) is broken at roughly 65 *k*_*B*_*T* nm^−2^ [26], which is consistent with the maximum longitudinal bending stiffness of the tropomyosin coiled-coil used in [27]. For short peptide chains, the elastic constant *k* is reduced to values around 40 *k*_*B*_*T* nm^−2^ [28]. The friction coefficient *γ* is estimated using the Stokes law *γ* = 6*πηR* with the dynamic viscosity of water at 300 K, *η*_*w*_ = 0.85 pN ns nm^−2^ [29], and assuming the amino acid radius *R* to be of the order of *l*_0_*/*2. We estimate a maximum strength of the hydrophobic interactions of 4 *k*_*B*_*T*, that corresponds to the hydration potential of isoleucine [30], although accurate measurements of the solvation free energies of biomolecules are difficult to access experimentally [31, 32]. Since the CC structure is not melting, but sliding, this value for *U*_0_ might be overestimating the hydrophobic interaction established between the residues of opposite strands. Similarly, the energy required for transferring buried residues from the interior to the protein surface is much lower than its full solvation [30]. Since the Rab5-EEA1 interaction and the consequent signal propagation occur without GTP hydrolysis [3], the activation energy *E*_*A*_ must be less than 20 *k*_*B*_*T*.

The ranges of *k* and *U*_0_ that would allow the signal to propagate until the opposite end of the EEA1 protein are shown in Fig. 3. The diffusion constant *D* has been obtained by fitting the solution to diffusion equation (Gaussian distribution) in an ubounded domain to the average of 2000 kink trajectories generated at the center of the chain. The diffusion constant *D* for kinks introduced both at the center and at the extreme of the chain are equivalent in the long time limit.

**FIG. 3.**
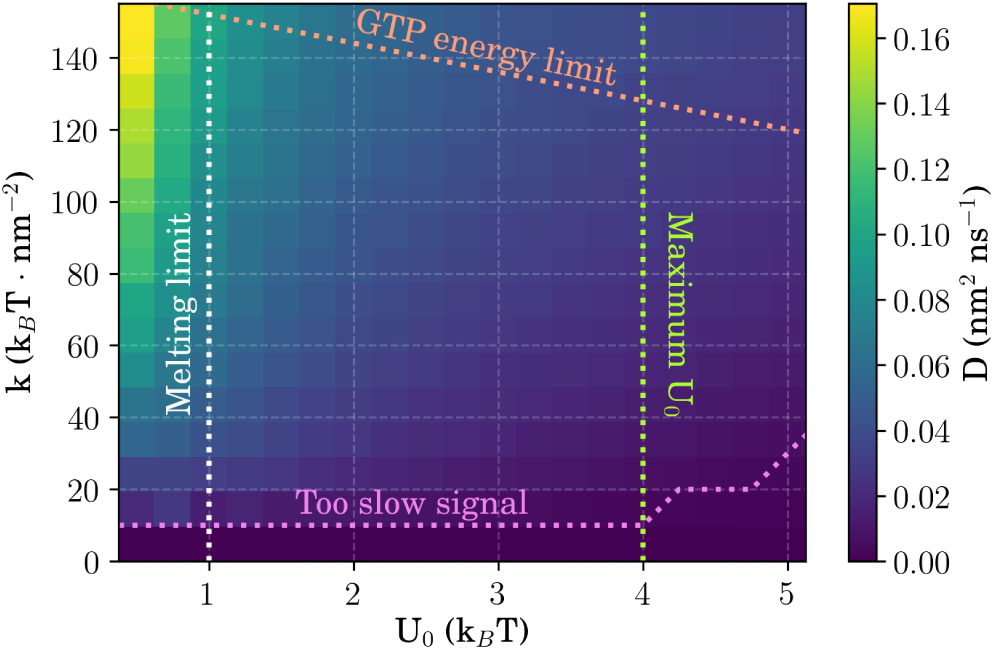
Diffusion coefficient of a kink generated in the center of a Frenkel-Kontorova chain for different pairs of *k* and *U*_0_, each extracted from the energy average of 2000 kink trajectories. The upper limit (orange) corresponds to the activation energy E_*A*_ required to overcome the potential energy barrier for energies below that released by GTP hydrolysis (*E*_*A*_ = 20 *k*_*B*_ *T*). The magenta line limits the cases where the diffusion constant is too low to be able to transmit the signal within the maximum allowed time, *t*_*c*_ = 0.1*s*. Hydrophobic energy differences *U*_0_ below 1 *k*_*B*_ *T* lead to unstable chains (white). Finally, the maximum energy released between a pair of hydrophobic residues *U*_0_ is about 4 *k*_*B*_ *T* [30] (shown in light green).

The sum of the energy contributions required to overcome the potential barrier *U*_0_, and to compress the first monomer to a length *l*_0_, must not surpass the energy released by the hydrolysis of a GTP molecule (≈ 20*k*_*B*_*T*),

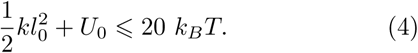

This provides an upper limit for the values of the elastic constant *k* and the potential well *U*_0_ (shown in orange). The time required to complete a full EEA1-Rab5 cycle is around *t*_*c*_ = 0.1 s [3]. This delimits the minimum required signal speed according to *t*_*c*_ ⩾ ⟨*N* ^2^⟩ */*2*D*, where *N* is the length of the chain. From the results shown in Fig. 3 and the physiological parameters in Table I, we predict the ranges for the hydrophobic interaction energy *U*_0_ ∈ (1.0, 4.0) *k*_*B*_*T* and an elasticity ranging from *k* ∈ [10, 150] *k*_*B*_*T* nm^−2^ to *k* ∈ [10, 125] *k*_*B*_*T* nm^−2^ depending on *U*_0_, allowing for signal propagation in EEA1 at physiological conditions. Hydrophobic contacts *U*_0_ with less than 1 *k*_*B*_*T* lead to unstable chains due to the spontaneous generation of kink-antikink pairs already observed in the results (white dotted line).

**TABLE I.** Choice of biologically relevant parameters for the Frenkel-Kontorova model applied to describe ideal coiled-coil structures.

| Parameter | Range | Reference |
| --- | --- | --- |
| Turn length, $L, l_0$ | 0.5 nm | [24, 25] |
| Chain length, $N$ | 400 beads | [33] |
| Hydrophobic int., $U_0$ | $< 4 k_B T$ | [30] |
| Elastic constant, $k$ | $1 - 65 k_B T \text{ nm}^{-2}$ | [26–28] |
| Dyn. water viscosity, $\eta_w$ | $0.85 \text{ pN ns nm}^{-2}$ | [29] |
| Activation energy, $E_A$ | $< 20 k_B T$ | [34] |

## d. Conclusions

How a local interaction affects the global property of a molecule is an important question in physical and biological systems. Recent experiments on the membrane tether EEA1 have shown that binding of the small GTPase Rab5 to the N-terminus of EEA1 significantly reduces the bending stiffness of this long coiled-coil protein. To elucidate this novel phenomenon, we proposed a simplified description of the CC as a finite length FK chain, modeling ligand binding as a register shift in the chain. Using numerical simulations, we found that the signal can reach the other end of the chain only in the presence of thermal fluctuations. Importantly, we determine the range of parameters that enable the intramolecular communication within the experimentally estimated time; those can be used to predict the elastic parameters and the hydrophobic interaction strength of the EEA1 CC.

Experimentally it has been found that the flexibility transition occurs only in the presence of discontinuities. Within our model, the discontinues can be considered as defects in the FK potential, which enable a wider fluctuation of FK monomers. Therefore, we can interpret the experimental finding as follows: in the presence of discontinuities, the binding of Rab5 can trigger a significant distortion of EEA1, which is revealed as a reduced bending stiffness. In the mutant EEA1 (without discontinuities), however, the signal can propagate through the whole chain while the chain remains stable. Our results indicate that as EEA1 remains in the vicinity of the critical point between stable and unstable state, it can dramatically change its global flexibility upon being triggered by the ligand. Including the discontinuities in our model is a future direction of research.

## ACKNOWLEDGMENTS

We acknowledge helpful discussions with M. Zerial and D. Murray. This research was supported by the Max Planck Society.

